# Production of MarathonRT and its comparison with commercial reverse transcriptases for tRNA sequencing library preparation

**DOI:** 10.64898/2026.03.09.710538

**Authors:** Jenni K. Pedor, Pavlina Gregorova, Matea Radesic, L. Peter Sarin

## Abstract

Over the past decade, groundbreaking discoveries have cemented transfer RNAs (tRNAs) as versatile regulators of translation and cellular function. As tRNA research gains momentum, several high-throughput sequencing methods have emerged for quantitative analysis of tRNA isoacceptors in cells. However, the strong secondary structure and rich post-transcriptional modification of most tRNA molecules pose significant challenges for reverse transcriptases, thus hampering library preparation and introducing quantification biases. Current approaches rely on processive next generation reverse transcriptases (ngRTs)—they successfully overcome these problems, albeit with the potential caveat of high experimental costs. Here, we introduce a recombinant MarathonRT (MRT) protein both with and without a C-terminal chitin binding domain (CBD), for which we present a simple and robust one-step purification protocol that yields over 26,000 enzymatic reactions per 0.5 L of expression culture. We also developed an affordable colorimetry-based method for determining the specific activity of these enzymes. Importantly, we benchmarked our in-house produced MRT and MRT-CBD enzymes using the mim-tRNAseq workflow and show that their performance match that of commercially available ngRTs. In addition, we implemented the use of a rapid tRNA spin column-based extraction method for tRNA-seq and LC-MS applications, establishing it as a viable alternative to conventional gel-extraction. Combined, this improved workflow significantly reduces the time and cost of tRNA-seq library preparation while providing an easily implementable MarathonRT purification protocol.

## Introduction

Transfer RNAs (tRNAs) are critical components in various essential biological processes, ranging from translation^1^, amino acid metabolism^2,3^, gene expression regulation (reviewed in ^4^), and priming of reverse transcription in retroviruses^5^ to biotechnological applications (reviewed in ^6,7^). Given their pervasiveness, sudden changes in tRNA abundance and function can lead to various diseases and dysfunctions (reviewed in ^7,8^). Hence, accurate quantification of tRNA is important not only for understanding cellular physiology and disease, but also for advancing tRNA-based therapeutics.

Over the past decade, significant efforts have been made towards developing tRNA quantification methods based on next-generation sequencing^9^. However, tRNA is notorious for its robust structure, high sequence similarity and extensive post-transcriptional modification (PTM) of its nucleosides. Based on their position, PTMs can be broadly categorized into two groups: tRNA body or anticodon stem loop modifications. PTMs located in the tRNA body are mostly associated with folding and stability, whereas modifications at the anticodon stem loop directly affect transcript decoding (reviewed in^10^). Next to PTMs, strong secondary structures and chemical modifications often create unsurmountable roadblocks for conventional reverse transcriptases (RTs), causing them to stall and dissociate from the template. This produces prematurely terminated reads that are difficult to map due to the high sequence conservation of tRNAs (reviewed in^8,9^). Taken together, these features make tRNAs one of the most technically challenging RNA molecules to sequence.

To overcome these challenges, numerous strategies including the enzymatic removal of common modifications^11–14^, limited alkaline hydrolysis^15^, and the use of novel group II intron encoded RTs^13,14,16,17^ have been implemented. Compared to conventional retroviral RTs, group II intron-encoded RTs are of retrotransposon origin and possess superior processivity on structured and/or modified templates^18,19^. Consequently, these next generation RTs (ngRTs) that include TGIRT^18^, MarathonRT (MRT)^19,20^, and recent commercial alternatives, such as uMRT (RNAConnect) and Induro (New England Biolabs, NEB), have become the RTs of choice in the latest Illumina-based tRNA-seq methods^14,16,17^. Importantly, ngRTs can produce full-length cDNA from challenging templates without enzymatic removal of modifications and without the need to destabilize the secondary structures by higher RT reaction temperature^14,16,17^. In addition, ngRTs are used in Oxford Nanopore Technologies (ONT) direct RNA sequencing applications to improve signal quality by destructuring the tRNA through cDNA synthesis, thus yielding a linearized tRNA molecule^21,22^. Currently, Induro is the officially recommended ngRT enzyme used in the ONT workflow.

Despite these advances, the availability of ngRTs and overall experimental costs remain significant bottlenecks in tRNA-seq applications. Commercial ngRTs offer high fidelity and processivity but can be prohibitively expensive for large-scale library preparations. Alternatively, researchers may opt to express recombinant MarathonRT (MRT) in-house using the Addgene-distributed plasmid (#109029). However, previously published protocols for expression and purification of MRT^19,20^ were developed for resolving the crystal structure of the MRT enzyme. As this seminal work required a very pure product, the resulting multi-step purification protocol was complex and laborious, heavily emphasizing MRT purity over yield and ease-of-production.

Similarly, in tRNA-seq workflows, a lot of time and effort is required to isolate bulk tRNA from total RNA. Denaturing polyacrylamide (PAA) gel-based extraction is the established albeit time-consuming and labour-intense method-of-choice in most protocols^14,16^. Alternative strategies include anion-exchange chromatography^23^ or reverse transcription of tRNAs using DNA splints^17^, which both have caveats. The former requires less manual labour but necessitate overnight precipitation steps and are not easily scalable, whereas the latter may be suitable for tRNA-seq, but does not work for mass spectrometry (MS)-based tRNA analysis. There are also commercial kits that capture short RNAs [e.g. miRNeasy Kit (Qiagen) < 200 nt or mirVana miRNA kit (Ambion) < 200 nt], but none of these have a cut-off that specifically enriches for tRNA without capturing 5S and 5.8S rRNA contaminants, which is not desired. However, a recently introduced silica-based spin column protocol enables rapid enrichment of tRNA from total RNA within minutes^24^, thus promising an efficient approach for preparing sequencing and MS-grade tRNA samples.

In this study, we present a simplified method for the expression and one-step purification of recombinant MRT and its C-terminally modified variant, MarathonRT-CBD (MRT-CBD). By inducing MRT expression at the early exponential phase, eliminating tag-cleavage, and removing non-essential purification steps, we obtain high yields of MRT with a performance comparable to commercial ngRTs at a fraction of the cost. In addition, removal of non-essential purification steps shortens the MRT production workflow by at least one day. Furthermore, we established rapid silica-based spin column tRNA enrichment method as an efficient alternative to gel purification and demonstrate its compatibility with tRNA-seq and LC-MS-based PTM profiling. The resulting workflow provides a cost-effective, reproducible, and accessible method for tRNA sequencing library preparation, facilitating broader accessibility of tRNA-seq technologies across molecular biology laboratories.

## Results

### An improved protocol for the expression and purification of MarathonRT

As most tRNA-seq protocols rely on highly processive ngRTs to convert tRNA to cDNA prior to sequencing library preparation, we set out to simplify the production process of MRT while making sure to retain the enzymatic activity. During our initial tests, we observed that inducing the expression of the protein at OD_600_ of 0.8 – 2^20,25^ persistently resulted in the majority of the protein precipitating during affinity purification. Secondly, the enzymatic removal of the N-terminal 6×His-SUMO-tag following affinity purification triggered excessive protein precipitation as described in the original publication^20^. Consequently, very little soluble protein remained available for subsequent chromatography steps, which ultimately resulted in a yield too small for downstream applications and an excessive cost-per-reaction. To resolve these issues, we first revised the expression constructs. In addition to the original MRT construct, we also introduced a Chitin Binding Domain (CBD) to the C-terminus of MRT (MRT-CBD) (Figure 1A), thereby “sandwiching” the protein with a SUMO and CBD tag to enhance protein solubility^26^. Secondly, we decided to omit the tag-cleavage step as another group II intron-encoded RT has been shown to function with the expression and solubility tags^18^. Thirdly, we also explored if using solely an affinity purification step yields a protein sample preparation with no competing enzymatic activity nor RNase contamination. The resulting expression and purification workflow is shown on Figure 1B. We observed that for both the MRT and the MRT-CBD construct, there was no precipitation during affinity purification when expression was induced in an *E. coli* Rosetta 2(DE3)pLysS strain at OD_600_ of 0.4 – 0.5 and grown for 18 h at 16 °C. We also noted that using transformants older than 3 days significantly reduces the protein yield and increases precipitation, which may be accounted for by mutations that accumulate in the coding gene. Following cell lysis by ultrasonication, we performed a one-step gravity flow affinity chromatography purification of the His-tagged proteins using a self-packed Ni-NTA column (1 mL resin/500 mL culture). The MRT and MRT-CBD containing elution fractions were pooled prior to buffer exchange and concentration. This yielded protein batches consisting primarily of full-length MRT or MRT-CBD along with some truncated protein species and other co-purifying native *E. coli* proteins (Figure 1C, D). The protein identities were confirmed by Western Blot, where it was clearly detectable that MRT-CBD expression yielded 25 % more of full-length product compared to MRT (Figure 1 D, Supp. File S1). Moreover, we consistently observed a specific set of truncated protein products that were clearly detectable by SDS-PAGE analysis and Western Blot using an anti-His-tag antibody (Figure 1D). We assume that these C-terminally truncated proteins are MRT cleavage products generated by *E. coli* proteases during the expression-purification protocol. Indeed, SDS-PAGE analysis revealed that these bands migrate faster following SUMO-tag cleavage (Supp. Fig. S1, A), which is consistent with a size-reduction following loss of SUMO-tag.

**Figure 1.**
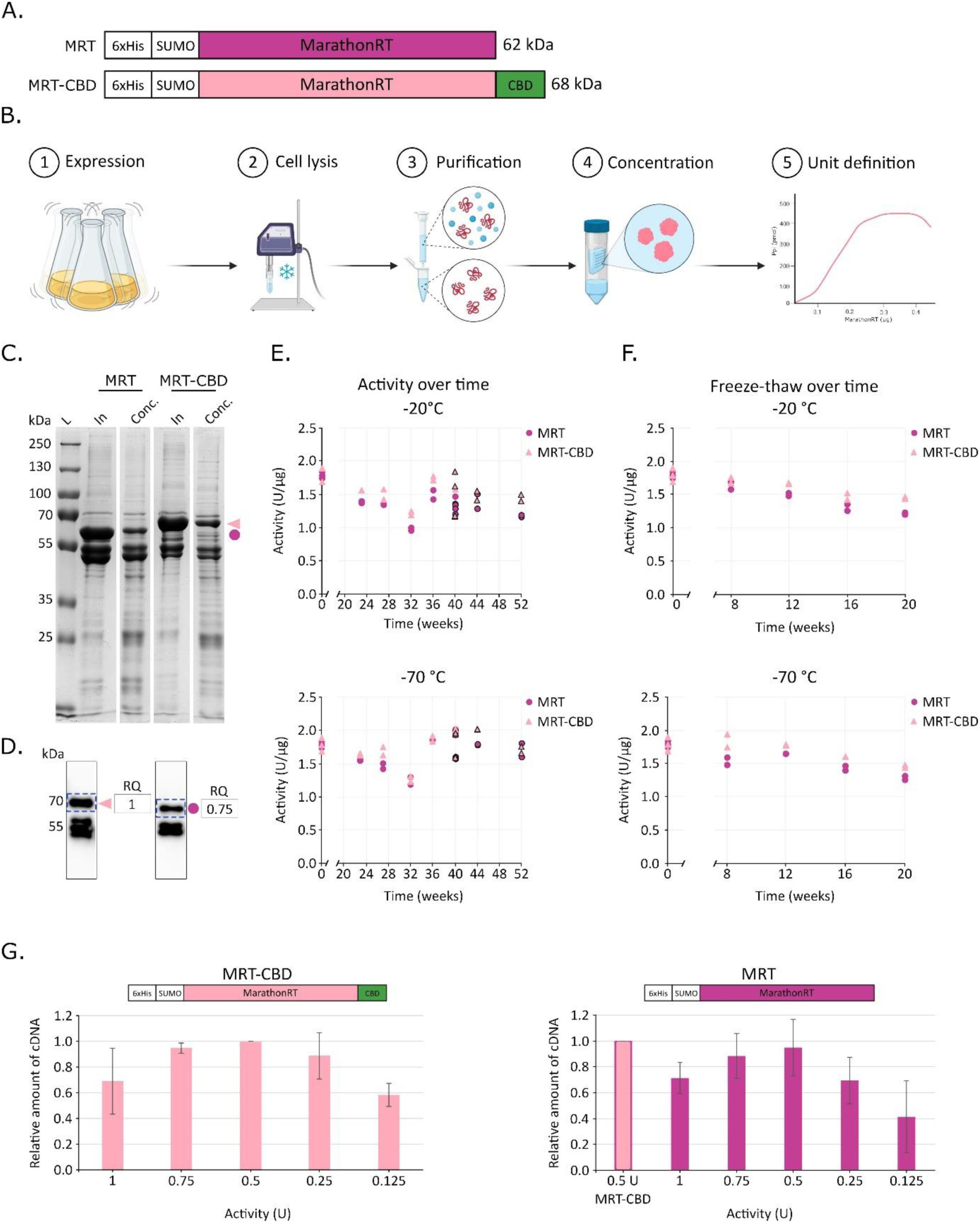
MRT and MRT-CBD construct design, purification, storability, and optimal enzyme-to-template ratio. (A) Schematic representation (not to scale) of MRT and updated MRT-CBD construct in a new plasmid backbone with a C-terminal CBD domain. (B) Overview of the optimized one-step protein production workflow. (C) Representative 10 % SDS-PAGE gel (1.5 mm). *In* – pooled elution fractions from affinity purification; *Conc* – concentrated protein. Bands representing full-length protein are indicated (dot = MRT, triangle = MRT-CBD) corresponding to panel A colours. Marker: PageRuler Broad Range Unstained Protein Ladder (Fisher Scientific). (D) Representative Western blot of concentrated protein detected by anti-His antibody. Full-length protein indicated by dot or triangle. Densitometry analysis using ImageLab v 6.0 volume box feature of the relative quantity (RQ) of the full-length protein shown next to the plot, full-length protein band marked by blue dashed line. (E, F) Protein storability. Top panels: activity of samples stored at −20 °C; bottom panels: activity of samples stored at −70 °C. Measurements were performed in technical duplicates, except the first time point (n = 3). Black outline around shapes indicates the use of a new RNA template batch. (E) Protein activity over 52 weeks. (F) Activity during repeated freeze-thaw cycles over 20 weeks. (G) Bar plot of densitometry analysis of free cDNA produced by indicated enzyme amount (U) from 100 ng of 3′-adapter-ligated tRNA template. 0.5 units of the same MRT-CBD reaction were loaded on both gels for comparison.

Consequently, using our revised fusion protein design along with the optimized production protocol, we obtained a total protein yield of 7.5 mg and 5.3 mg per 0.5 L of culture for MRT and MRT-CBD, respectively (Table1; Supp. File S2). Furthermore, we did not detect any precipitate at any point of the purification process, which suggests that our protocol significantly mitigates MRT precipitation and thus increases the yield of the active enzyme.

### MRT and MRT-CBD retain their enzymatic activity over a 12-month storage period

To quantitatively determine the activity of our in-house produced MRTs and normalize against batch-to-batch variation, we adapted a simple and cost-effective colorimetric assay that measures pyrophosphate (PPi) release during the cDNA polymerization reaction (modified from^27,28^). In this assay, 1 enzyme unit (U) is defined as the amount of MRT/MRT-CBD needed to release 1 nmol of PPi in 30 min at 42 °C when using a defined RNA fragment as template and a gene-specific reverse transcription primer at standard reaction conditions^29^. Hence, the average activity of MRT and MRT-CBD protein preparation measured directly after purification was 1.79 ± 0.05 U/µg and 1.79 ± 0.11 U/µg, respectively, which equals a total of 13,478 U and 9,420 U per 0.5 L of culture, respectively (Table 1, Supp. File S3). Consequently, if 0.5 U is used for each standard RT reaction, this corresponds to 26,957 MRT and 18,841 MRT-CBD reactions, which results in a total cost-per-reaction of 0.0023€ or 0.0034€, respectively (Supp. File S4; prices correct at time of writing). Overall, the colorimetric assay demonstrated that our optimized expression and purification protocol yields enzymatically active RTs that can be affordably produced.

**Table 1.** MRT and MRT-CBD yield and production cost compared to commercial enzymes.

| **Protein** | **Total yield**<br>**(mg)** | **Activity**<br>**(U/mg)** | **Total units**<br>**(U)** | **Reactions**<br>**(nr)** | **Cost per rxn**<br>**(€)** |
| --- | --- | --- | --- | --- | --- |
| **MRT** | 7.5 | 1,790 | 13,478 | 26,957 | 0.0023 |
| **MRT-CBD** | 5.3 | 1,790 | 9,420 | 18,841 | 0.0034 |
| **uMRT**<br>**(R1002S)** | N/A | N/A | N/A | 20* | 19.7 |
| **Induro**<br>**(M0681L)** | N/A | N/A | N/A | 50* | 11.9 |

\*As tested, also available in other amounts.

Next, using the same colorimetric assay, we set out to monitor the long-term stability and storability of the MRT (6.3 mg/mL) and MRT-CBD (4.4 mg/mL) protein preparation stocks (Supp. File S2) when stored at −20 °C or −70 °C for up to 12 months (Figure 1 E, Supp. File S3). As previously established, the initial activity for MRT and MRT-CBD was 1.79 ± 0.05 and U/µg 1.79 ± 0.11 U/µg, respectively, which after 12 months of continuous storage at −20 °C decreased to 1.18 ± 0.02 U/µg and 1.45 ± 0.07 U/µg, respectively (Figure 1 E). This suggests that MRT-CBD is marginally more stable and retains a higher specific activity at −20 °C. Conversely, we observed no significant loss of activity for either MRT or MRT-CBD when stored at −70 °C for 12 months, as both enzymes featured an average activity of 1.71 ± 0.14 U/µg and 1.71 ± 0.07 U/µg (Figure 1 E, Supp. File S3), indicating that cryogenic conditions are preferential for maintaining enzymatic activity during long-term storage. However, we noticed a significant decrease in activity for both enzymes and storage conditions after 32 weeks (∼7 months). Since this change was sudden and both conditions were equally affected, we suspected a potential issue with the activity assay, possibly due to degradation of the RNA template. Consequently, a new batch of the template (Figure 1E; data points with black outline) was produced and assayed in parallel with the old batch (Figure 1E; data points without any outline) at weeks 36 and 40. This revealed that the old and new template batches produced very similar results and that the average specific activities obtained for both enzymes and storage conditions were equivalent to the initial values at week 0, suggesting that the reduction of activity observed at week 32 can most likely be accounted to a technical mishap.

In addition to continuous long-term storage, we also assessed how multiple freeze-thaw cycles affected the stability of both MRT enzymes. To this end, we measured MRT and MRT-CBD activity following 5 freeze-thaw cycles of the same enzyme samples spanning 20 weeks of storage at −20 °C and −70 °C, respectively. Unsurprisingly, each subsequent freeze-thaw cycle caused a loss of activity irrespective of enzyme or storage condition, although enzymes stored at −70 °C performed marginally better (Figure 1F). At the end of the storage period, MRT activity had dropped from an initial average of 1.79 ± 0.05 U/µg to 1.21 ± 0.02 U/µg or 1.28 ± 0.03 U/µg, respectively, when stored at −20 °C or −70 °C. However, MRT-CBD was less affected, possibly due to the protection offered by the C-terminal CBD-tag, and we observed a loss of activity from an initial average of 1.79 ± 0.11 U/µg to 1.45 ± 0.02 U/µg or 1.46 ± 0.03 U/µg, respectively, when stored at −20 °C or −70 °C (Figure 1F, Supp. File S5). Finally, high protein concentrations have been reported to act as cryoprotectants and stabilizers against freeze-thaw induced degradation^30^, albeit such protective properties are lost upon dilution. To establish the stability of MRT solutions diluted to 0.6 µg/µL, which corresponds to ∼1.5 U/µL, we exposed these to 2 or 4 freeze-thaw cycles spanning 7 or 16 weeks at −20 °C or −70 °C, respectively (Supp. Fig. S2A; Supp. File S6). At these conditions, we noted a small activity drop for both MRT and MRT-CBD, although as with the concentrated enzyme stocks, enzymes stored at −70 °C yielded higher activities (Supp. Fig. S2A; Supp. File S6).

### Excess MRT and MRT-CBD reduces overall cDNA production efficiency

After determining the specific activity for the enzymes, we defined the optimal amount of enzyme needed for processing 100 ng of tRNA, as this constitutes the input amount for the RT reaction in the mim-tRNAseq protocol^31^. Hence, we systematically performed RT reactions using 0.125-1.75 U of MRT or MRT-CBD and the resulting cDNA formation was characterized by urea-PAA gel electrophoresis and subsequent image densitometry analysis (Supp. Fig. S2, C; Supp. File S7). This revealed that the highest amount of cDNA was obtained with 0.5 U of MRT or MRT-CBD in the reaction (Figure 1G). Importantly, our analyses also showed that additional MRT or MRT-CBD (>0.5 U for 100 ng of tRNA) did not improve cDNA synthesis—on the contrary, an excess of enzyme (0.75-1.75 U) induced the formation of high molecular weight complexes consisting of MRT or MRT-CBD invariantly bound to the synthesized cDNA, which lowered the yield of available cDNA (Supp. Fig. S2C). MRT is known to have a highly basic secondary RNA binding site, which can contribute to the higher binding rate of the template tRNA and render the complex inactive^19^. Consequently, the most optimal activity for MRT or MRT-CBD in the tRNA sequencing workflow is achieved when using 0.5 U of enzyme per 100 ng of tRNA template.

### Enhanced tRNA enrichment by silica-based spin columns facilitates sample preparation

The first step in tRNA-seq and MS-based tRNA PTM detection is to obtain high-quality tRNA. A recently published protocol for tRNA enrichment using silica-based spin columns^24^ offers a potential alternative to the established methods^23^. In this protocol, tRNAs and other short RNAs (<120 nt) are enriched from total RNA by a simple four-step process that takes approx. 30 min (Figure 2A). Importantly, this approach does not enrich for 5S or 5.8S rRNA, which are common contaminants in many RNA isolation procedures. The original protocol uses Macherey-Nagel NucleoSpin RNA columns, where 80-100 µg total RNA is needed for efficient tRNA extraction. However, this is an excessively high amount for tRNA sequencing, where only a fraction of the material is needed for cDNA synthesis. Therefore, we assessed the protocol for use with silica-based spin columns that are optimized for low sample amounts and volumes, such as NucleoSpin RNA XS (Macherey-Nagel) and Monarch Spin RNA Cleanup (10 µg) (NEB) columns. Interestingly, the NucleoSpin RNA XS column did not achieve tRNA and rRNA separation even at total RNA input amounts as low as 0.75-2.0 µg (Supp. Fig. S3A). The Monarch® Spin RNA Cleanup (10 µg) column proved to be the better choice as it performed similarly to the original protocol and efficiently enriched for tRNA at the tested total RNA inputs of 2.5 µg, 5 µg, or 10 µg (Figure 2 B). Furthermore, we show that the cut-off for long RNA binding (Figure 2A, step I) to the Monarch Spin RNA Cleanup (10 µg) column can be modulated by adjusting the EGDA concentration, where 35 % EGDA provides the best balance between tRNA enrichment and rRNA removal (Supp. Fig. S3B) as used in the original protocol^24^. Nonetheless, we did observe that a small fraction of the tRNA remains bound to the first column (Figure 2C), which constitutes a minor trade-off in favour of enhanced separation. To benchmark these adaptations, we applied the same starting material (total RNA from *S. cerevisiae*) for both tRNA gel extraction and spin column-based tRNA enrichment (Figure 2D). As expected, we achieved a recovery rate of 48 % for the tRNA fraction using the column-based method, which is slightly higher than the 37 % yield obtained with gel extraction (Table 3). We also noted the faint presence of 5S rRNA in both column and gel enriched samples (Figure 2D), which was attributed to extraction-dependent differences in total RNA composition (Supplementary Results 1, Supp. Fig. S3D, E).

**Figure 2.**
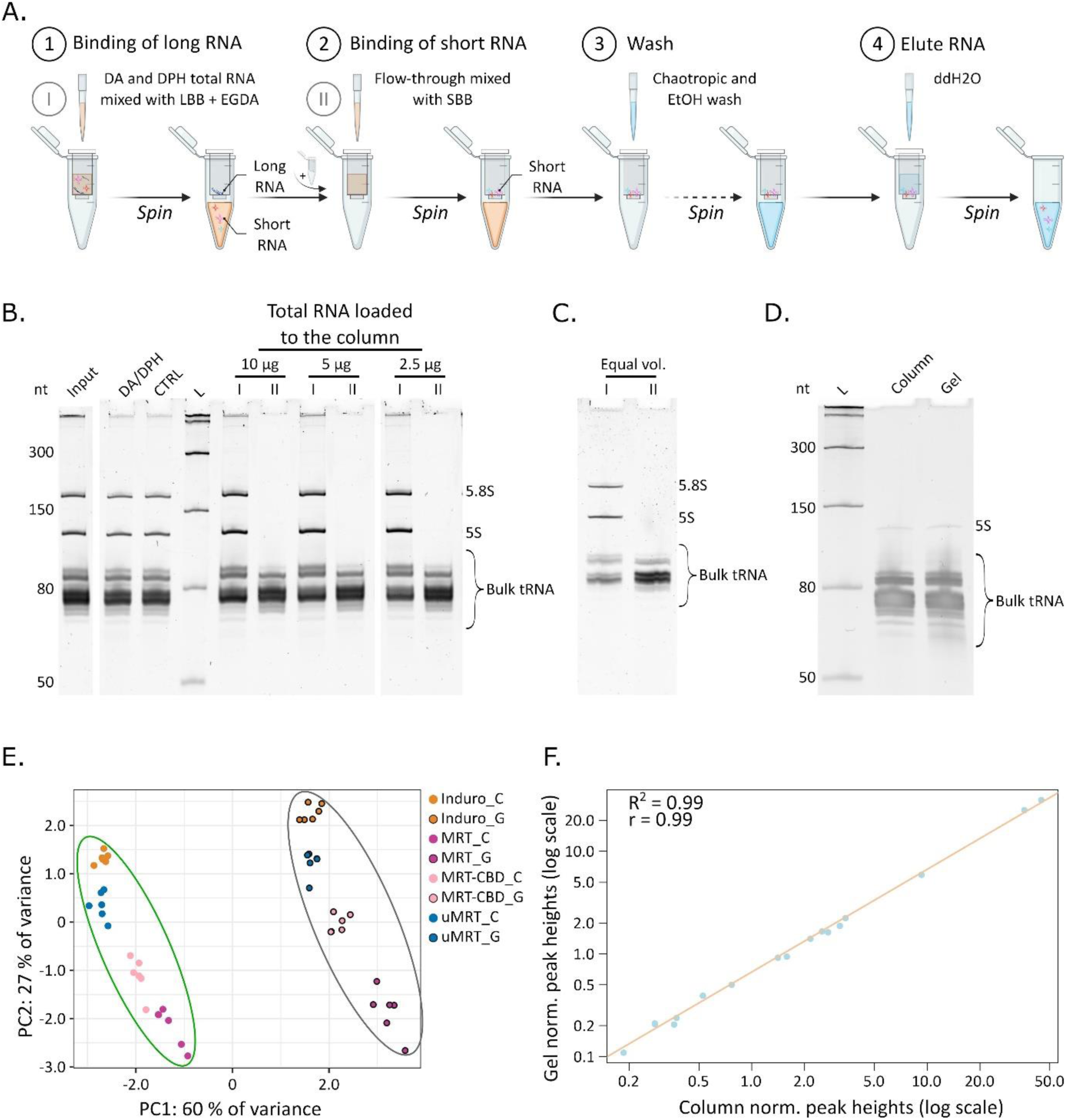
tRNA column and gel extraction for sequencing. (A) Schematic overview of the tRNA silica-based spin column purification process. First, the total RNA sample is mixed with long RNA binding buffer (LBB) and bound to the first column. The flowthrough will contain RNAs < 120 bp. This flowthrough is mixed with short RNA binding buffer (SBB) and bound to the second column. The RNA bound to the column is then washed with a chaotropic wash (CW), ethanol wash (EW) and eluted with ultra-pure water. (B–D) RNA samples analysed on 8–10 % denaturing urea-PAA gels. Marker: Low Range ssRNA (NEB). Bands corresponding to 5.8S and 5S rRNA are indicated. (B) Binding capacity of Monarch RNA Cleanup Columns (10 μg, NEB) loaded with 10, 5, or 2.5 µg total RNA. 50 ng RNA loaded to gel from column I and II eluates. *Input* – total RNA post-phenol/BCP extraction; *DA/DPH* – deacylated and dephosphorylated total RNA; *CTRL* – only dephosphorylated total RNA; *I/II* – eluates from column I and II. (C) Equal volumes loaded on gel from column I and II eluates. (D) Column and gel-extracted tRNA used in the tRNA-seq library preparation. (E) PCA clustering of sequenced samples (n = 48). Green circle: column extracted tRNA; black circle: gel-extracted tRNA. (F) Correlation of normalized peak heights of global tRNA modifications detected by LC-MS between extraction methods (log_2_ scale; column extraction n = 5, gel extraction n=3). Pearson’s *r* = 0.9998, R² = 0.9995.

Although our samples contain traces of rRNA, previous tRNA sequencing studies have successfully used samples containing RNAs shorter than 200 nt (incl. rRNA)^13^. Consequently, we proceeded with the downstream analysis.

### Choice of tRNA enrichment method does not bias tRNA-seq or MS analyses

To validate our approach, we generated sequencing libraries from gel-extracted and column-enriched tRNA samples using our in-house produced MRT and MRT-CBD and benchmarking them against Induro and uMRT (Figure 3A). We used total RNA from *S. cerevisiae* and followed the mim-tRNAseq protocol^31^. Briefly, we generated 8 libraries, each consisting of 6 technical replicates per ngRT and tRNA extraction method, which were pooled into 1 equimolar sequencing library and sequenced on Illumina NovaSeqX platform (Supplementary results 2, Supp. Fig. S4-S6). Illumina sequencing generated 1.07 billion reads, with 73 % assigned and 27 % unassigned. Over 88 % of bases had the quality score ≥ Q30 (Supp. File S8). After the analysis of R1 reads using the mim-tRNAseq toolkit^31^ we identified that all 8 libraries had the proportion of uniquely mapped reads ≥ 97 % (Supp. Fig. S7, A; Supp. File S8). Following the principal component analysis of the sequenced samples, we observed that samples extracted using silica-based spin columns clustered together, separating clearly from the cluster formed by the gel-extracted samples (Figure 2E). Importantly, all detectable 42 *S. cerevisiae* cytosolic tRNA isoacceptors^32^ (Supp. File S9) that were identified could be found to an equal extent in both spin-column and gel-extracted samples with read count correlation *r* = 0.93-0.96 (Supp. Fig. S8). To verify that column enrichment does not negatively affect the chemical modifications present on tRNAs, we performed LC-MS analyses of tRNA samples from column-enriched and gel-extracted samples. We obtained a Pearson’s correlation of 0.99 between the samples with most of the detected ions corresponding to canonical nucleosides and PTMs found on tRNA (Figure 2F, Supp. File S10), indicating the very similar performance of these two extraction methods. However, we did identify traces of m^6^_2_A in two of the gel-extracted samples, but not in any of the column-enriched samples (Supp. File S10). Since m^6^_2_A is a rRNA-specific modification^33^, it suggests that some of the gel-extracted samples contained a minor rRNA contamination. Overall, our results show that the performance of the silica-based spin column tRNA enrichment method is comparable to or even slightly exceeds established gel extraction-based approaches.

**Figure 3.**
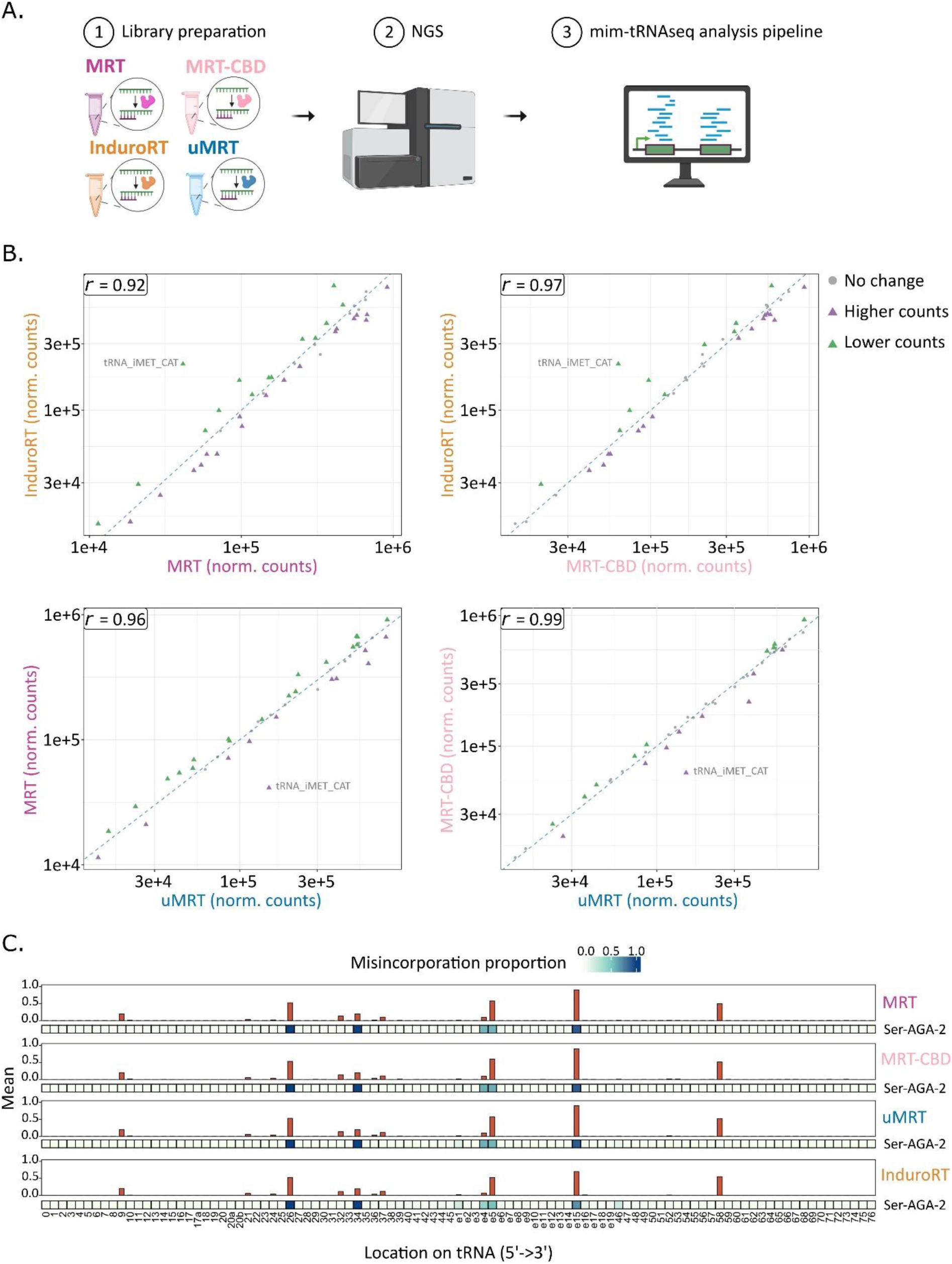
Benchmarking in-house MRT and MRT-CBD versus commercial RT enzymes. (A) Schematic overview of the tRNA-seq workflow. (B) Reproducibility of tRNA quantification across RT enzymes. Scatter plots show pairwise comparisons of normalized tRNA counts (technical replicates n = 6). Axes represent normalized read counts from DESeq2 (adj-p ≤ 0.05). Pearson (*r*) indicated. (C) Mean position-specific mismatch frequencies across tRNAs aggregated over technical replicates (n = 6) per enzyme. Only positions with coverage >0.0005 are shown. Misincorporation pattern for tRNA-Ser-AGA-2 is shown under each enzyme.

### MRT and MRT-CBD performance is on par with uMRT and Induro

Next, we proceeded with comparing the performance of our in-house produced ngRTs with the commercial enzymes using the sequencing results obtained from the above generated libraries where column-enriched tRNA was used as starting material. Unexpectedly, our analysis of the detected cytosolic tRNA species (clustered by anticodon) counts showed that tRNAs reverse transcribed by MRT-CBD clustered closer to uMRT and Induro than to MRT (Supp. Fig. S7, B). Indeed, we also observed that MRT-CBD/uMRT and MRT-CBD/Induro samples yielded a high Pearson’s correlation with *r* = 0.99 and *r* = 0.97, respectively (Figure 3B), whereas Induro/MRT samples varied the most with a correlation of *r* = 0.92 and the remaining samples showed an intermediate correlation of *r* = 0.96-0.98 (Figure 3B; Supp. Fig. S7C). This suggests that MRT-CBD yielded results that are similar to those obtained with Induro and uMRT, whereas MRT appears marginally divergent.

A closer inspection yielded some interesting observations for all ngRTs studied. First, samples reverse transcribed by Induro systematically showed higher initiator tRNA-Met-CAT counts than those generated by MRT-derived enzymes (Figure 3B; Supp. Fig. S7, C). Moreover, a similar overrepresentation was not observed for any other tRNA species. Second, the misincorporation pattern for these group II intron-encoded RTs is highly similar (Figure 3C). Positions 9, 26, e15 (refers to the extended variable loop as numbered in^34^) and 58 show consistent misincorporation at a high rate, whereas anticodon stem loop positions 32, 34, 36 and 37 show a slightly less frequent, but nonetheless distinguishable misincorporation pattern (Figure 3C). These coincide with frequent post-transcriptional modification of these sites in *S. cerevisiae* cytosolic RNA (reviewed in^35^)—with the notable exception of misincorporation at the variable loop present only for tRNA-Ser. This observation is unexpected as tRNA-Ser lacks a known modification at position e15^33^. Finally, the reverse transcription stop-rate remains low for all ngRTs (Supp. Fig. S7, D). MRT and MRT-CBD have a slightly higher 3’ bias compared to uMRT and Induro, whereas Induro shows the lowest 3’ bias (Supp. Fig. S9). In conclusion, our tRNA-seq results suggest that MRT and MRT-CBD enzyme preparations performed at a highly similar level to the commercial uMRT and Induro enzymes.

## Discussion

In this study, we have developed a simple and robust workflow for in-house production and purification of highly active MRT and MRT-CBD, supplemented with an easy-to-implement colorimetric assay for batch comparison and storability assessment (Figure 1). By optimizing construct design, inducing expression at lower culture density, and omitting protein-tag cleavage and additional chromatography steps (Figure 1B), this revised protocol consistently yields milligram amounts of soluble proteins from 500 mL of *E. coli* culture (Table 1). The resulting MRT and MRT-CBD preparations are stable over extended storage time and withstand multiple freeze–thaw cycles (Figure 1E, F; Supp. Fig. S2A). The increased fraction of full-length MRT-CBD and reduced accumulation of truncated species (Figure 1C, D) further underscores the stabilizing effect of “sandwiching” the enzyme between solubility tags^26^, in line with previous observations for other group II intron RTs^18^. Furthermore, the tags might act as replacement for natively bound RNA, which is necessary for group II intron RT stability^18^. Importantly, the performance of the ngRTs is comparable to that of the commercial Induro and uMRT, whilst being achieved at a fraction of the cost-per-reaction (0.0023–0.0034 €, excluding personnel costs) (Table 1).

To establish in-house produced MRT and MRT-CBD preparations as viable alternatives for tRNA-seq studies, we benchmarked their performance against commercial Induro and uMRT using the mim-tRNAseq workflow. This revealed that the in-house produced ngRTs support high-quality tRNA libraries with mapping rates, tRNA isoacceptor counts, misincorporation patterns, and RT stop profiles comparable to commercial enzymes (Figure 3; Supp. Fig. S7). MRT-CBD closely matches cytosolic tRNA isoacceptor counts obtained with Induro and uMRT while maintaining efficient readthrough at heavily modified positions, indicating that the extra CBD-tag does not adversely affect enzyme function. Minor differences, such as slightly lower tRNA_iMet_CAT counts in MRT libraries and higher tRNA_iMet_CAT counts in Induro libraries (Figure 3B; Supp. Fig. S7C), as well as a modest variation in 3’ bias (with Induro exhibiting the lowest 3’ bias) (Supp. Fig. S9), likely reflect subtle processivity differences rather than fundamental performance gaps. Importantly, the co-purified proteins present in the MRT and MRT-CBD protein preparations do not seem to have a negative effect on the enzyme, neither in terms of activity, specificity, or processivity. Overall, our results are consistent with previous studies and further highlight the comparable performance of group II intron-encoded RTs, as recently shown for Induro and TGIRT^17^. Our work further emphasizes the superior capabilities of ngRTs to provide essential information on tRNA sequence, PTMs and their location without any prior treatment of the template RNA^16,17^ (Figure 3, Supp. Fig. S7). Despite these advances, there is still room for further optimization in tRNA sequencing library preparation—for example, adding a periodate oxidation step enables the charging levels of tRNAs to be determined^36^, whereas applying the template-switching function of MRT eliminates the 3’adapter ligation step from the library preparation workflow^37^.

The one-step Ni-NTA protocol applied here yields enzymatically active MRT and MRT-CBD preparations devoid of competing enzymatic activities. Although this is sufficient for tRNA-seq, the presence of co-purifying bacterial proteins and truncated MRT species may limit their use in assays that demand very high purity, such as structural studies, single-molecule biophysics, or highly sensitive diagnostic assays. In such applications, additional purification steps, such as gel filtration or ion-exchange chromatography as performed in the original protocol^19,20,25^ may be required. In parallel, this study demonstrates that silica-based spin column tRNA extraction^24^ provides a time-efficient and scalable alternative to traditional denaturing gel-based tRNA extraction (Figure 2). The optimized two-step spin column protocol permits reliable enrichment of bulk tRNA from as little as 2.5 µg of total RNA, achieving average yields of 48 % compared to 37 % for gel extraction (Table 2). While we observed some 5S rRNA contamination due to the sensitivity to the phenol source in upstream total RNA extraction, these issues did not impair downstream tRNA-seq performance, as indicated by ≥97 % uniquely mapped reads in all libraries (Supp. Fig. S7A, Supp. File S8). Importantly, LC-MS analysis shows no detectable differences in PTM profiles between column- and gel-extracted tRNAs (Figure 2F, Supp. File S10), and tRNA-seq demonstrates that all identified tRNA species are present in both sample types across all tested RT enzymes (Supp. Fig. S8, Supp. File S9).

**Table 2.**
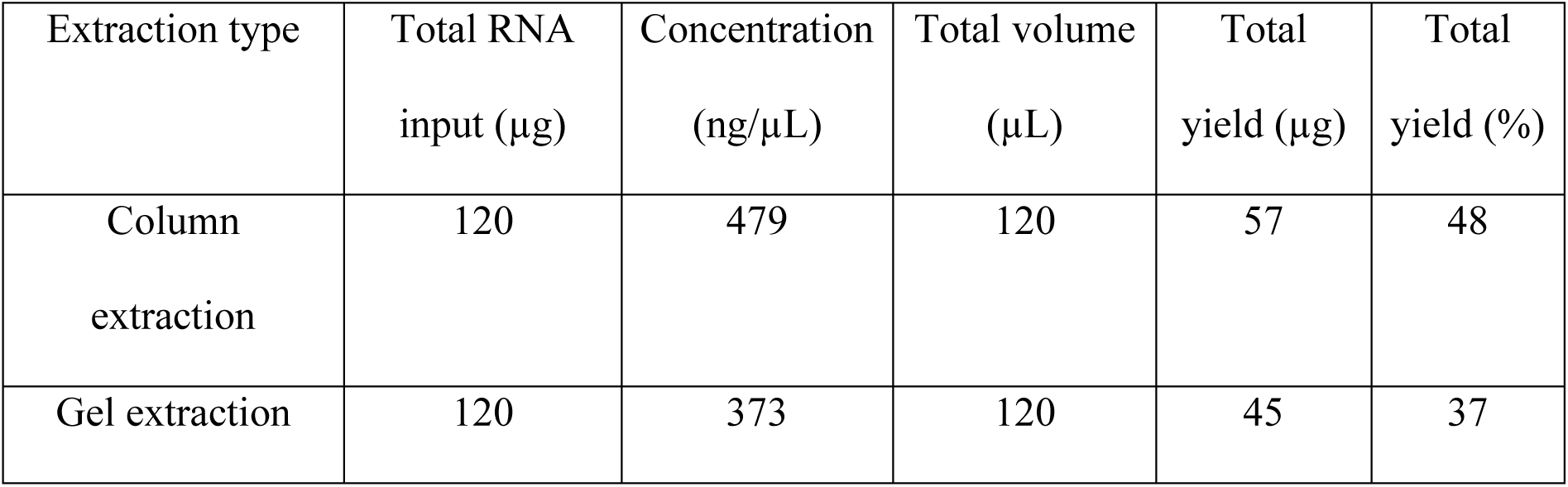
Comparison of tRNA enrichment efficiency by extraction method.

Although each isolation method has inherent biases—column enrichment may underrepresent the longest tRNAs near the membrane cut-off, whereas gel extraction may discriminate against slowly diffusing tRNA species—the overall isoacceptor profiles are highly similar, and both methods support quantitative tRNA profiling. Crucially, column-based enrichment for 6–12 samples can be completed in ∼30 minutes at room temperature, whereas conventional gel extraction requires 1–2 days and substantially more consumables, equipment, and hands-on time. In addition, we successfully used the modified one-step spin column-based RNA purification for *in vitro* transcribed template RNA directly from the reaction mixture^29^ (Material and Methods) and for tRNA purification after chemical treatments, such as de-acylation and de-phosphorylation^38^.

The combined MRT purification and spin column based tRNA extraction workflow thus lowers both the financial and technical barriers for implementing state-of-the-art-tRNA-seq and LC–MS-based tRNA modification analysis. All steps rely on standard molecular biology reagents and equipment^29^, and the enzyme expression plasmid is openly available, enabling straightforward adoption and further adaptation to different organisms, sample types, and tRNA-focused sequencing strategies. In addition, this work contributes to the further improvement of MRT and allows for more control over experimental design. Beyond tRNA-seq, the availability of highly active, low-cost in-house MRT and MRT-CBD opens the door for broader deployment of group II intron RTs in various applications, including RT-PCR on structured templates^39^, RNA structure probing^25^, modification mapping^40^, direct RNA and long-read sequencing^21,22^, and single-cell-seq^41^. In conclusion, our simplified and affordable MarathonRT production and purification protocol combined with silica-based column extraction of tRNA forms a robust, flexible, and accessible platform for advanced studies of tRNA biology and protein synthesis regulation.

## Materials and methods

### Bacterial strains and plasmids for cloning

pLJSRSF7-T7-6xHis-SUMO-MarathonRT-CBD plasmid (Addgene #253350) was constructed by amplifying MarathonRT from pET-6xHis-SUMO-MarathonRT plasmid gifted by Anna Pyle (Addgene plasmid #109029; originally described in^19^) using forward (5’-CATAGGATCCACATCATCATCATCATCACGGC-3’) and reverse (5’-ATCAGGTACCTCCGCAGGTCACGCATTTTTCGA-3’) primer (Metabion). The empty backbone pLJSRSF7 was gifted by Hideo Iwai (Addgene plasmid #64693; originally described in^42^) and was modified by deleting restriction fragment spanning from 85 bp (NcoI) to 1695 bp (BamHI). The amplified MRT fragment was inserted into modified pLJSRSF7 using BamHI and KpnI restriction sites. The resulting plasmids were amplified in *E*. *coli* DH5α.

### One-step purification of MRT-RT

MarathonRT purification protocol was modified from previous publications^19,20,25^. pLJSRSF7-T7-6xHis-SUMO-MarathonRT-CBD plasmid was transformed into *E. coli* strain Rosetta 2(DE3)pLysS competent cells. The cells were grown at 37 °C to an OD_600_ of 0.4, and recombinant protein expression was induced with 0.5 mM IPTG overnight (18 h) at 16 °C. The pellet from 500 mL culture was collected by centrifugation at 5,000 g for 15 min at 4 °C and washed with ice-cold 1× PBS followed by resuspension in buffer A (25 mM Na-HEPES, pH 7.5, 1 M NaCl, 10 % glycerol, 2 mM β-Mercaptoethanol) on ice. The suspension was sonicated for 5 min (30” on, 30” off) using a probe sonicator (Hielscher Ultrasonics), at 60 % amplitude and 0.5 cycle. The sample was centrifuged at 22,000 g for 45 min at 4 °C. The supernatant was filtered through 0.2 µm PES filter and incubated with Ni-NTA Resin (Thermo Fischer Scientific, 2 mL of 50 % slurry for 500 mL of bacterial culture) that had been pre-equilibrated in buffer A. After 1 h of incubation at 4 °C on an end-over-end shaker, the Ni-NTA resin-sample mixture was loaded onto a self-packable column. The resin was then washed with ice cold buffer A, and subsequently buffer B (25 mM Na-HEPES, pH 7.5, 500 mM NaCl, 10 % glycerol, 2 mM β-Mercaptoethanol, 20 mM imidazole), and C (25 mM Na-HEPES, pH 7.5, 500 mM NaCl, 10 % glycerol, 2 mM β-Mercaptoethanol, 30 mM imidazole). The protein was eluted with buffer D (25 mM Na-HEPES, pH 7.5, 300 mM NaCl, 10 % glycerol, 2 mM β-Mercaptoethanol, 300 mM imidazole) and collected by 1 mL fractions. The protein containing fractions were pooled and buffer exchanged for buffer G (25 mM K-HEPES, pH 7.5, 300 mM KCl, 50 % glycerol, 1 mM DTT) and concentrated into 1,200 µL using a centrifugal filter unit (Amicon Ultra Centrifugal Filter, 50 kDa MWCO, Merck). The concentrated protein was aliquoted and stored at −80 °C. Purified protein yield was quantified using Bradford assay^43^ and its purity assessed on SDS-PAGE with the expected size of 62 kDa for MRT and 68 kDa for MRT-CBD. For a more detailed purification protocol refer to^29^.

### Unit definition

#### RNA template preparation

Plasmid T7-CMVtrans-FFLuc-polyA gifted by Marcel Bruchez (Addgene plasmid #101156, originally described in^44^) was linearized in 40 µL reaction mixture containing 10 µg of the purified plasmid, 2 µL EcoRI (Thermo Fisher Scientific), 4 µL 10× Fast Digest Buffer and ddH_2_O up to 40 µL. The reaction was incubated for 1 h at 37 °C followed by inactivation of the enzyme at 80 °C for 5 min. The linearized plasmid was purified using NucleoSpin Gel and PCR Clean-up kit (Macherey-Nagel) following the manufacturer’s protocol and eluted into ddH_2_O. The template RNA was synthetized *in vitro* using T7 RNA polymerase (Thermo Fisher Scientific) in 50 µL transcription run-off reaction following the manufacturer’s protocol with slight modifications. In brief, we used 0.8 U of RNAsin Plus (Promega) and incubated the reaction for 2 h at 37 °C. Next, the DNA template was degraded by adding 2 µL of RQ1 RNase-Free DNase (Promega) and incubated for 15 min at 37 °C. The precipitated inorganic phosphate was removed by centrifugation at 16,000 g for 5 min at RT and the cleared supernatant was transferred into a clean microfuge tube. The RNA was purified on NucleoSpin RNA Columns for RNA purification (Macherey-Nagel, 740955.250S) using only the one-step purification method for products longer than 120 nt from^24^. The RNA (642 nt) concentration was measured by Qubit™ RNA BR Assay (Thermo Fisher Scientific) on a fluorometer (DeNovix) and purity assessed by denaturing electrophoresis on 5 % PAA/7 M urea/0.5× TBE gel. For a more detailed protocol refer to^29^.

#### Unit definition/Activity measurement

MRT/MRT-CBD activity was estimated by a colorimetric assay based on malachite green method for phosphate determination^27,28^. All the reagents were prepared/stored in plastic or glassware thoroughly rinsed with ddH_2_O to avoid any phosphate contamination. For a standard curve, inorganic phosphate (Pi, NaH_2_PO_4,_ Fisher BioReagents) dilutions ranging from 0 – 4,000 pmol were prepared from a fresh dilution of 50 µM Pi and ddH_2_O and added onto a microplate in triplicate. See more detailed protocol in^29^.

For 100 reactions, the detection solution (DS) was freshly prepared by mixing 1,574 µL malachite green reagent (0.12 % malachite green in 3 M sulfuric acid), 394 µL 7.5 % ammonium molybdate (Acros Organics) and 32 µL 10 % Tween in this specific order to avoid precipitation. For detection, 20 µL of DS solution was added to the Pi dilutions and incubated for 30 min at RT. The absorbance was measured at 620 nm using a Multiskan FC Microplate Photometer (Thermo Fisher Scientific). Next, to assess the activity of MRT/MRT-CBD, a reverse transcription reaction was performed in 20 µL final volume, where 300 ng of template RNA (described above) and 0.05 µM gene specific reverse transcription primer (5’-TCACTGCATACGACGATTCTG-3’) (Metabion) were mixed, incubated at 95 °C for 30 s, annealed for 5 min at RT and combined on ice with 1× MRT reaction buffer (50 mM Tris pH 8.3, 200 mM KCl, 2.5 mM MgCl_2_, 5 mM DTT, 20 % glycerol), 0.5 mM each dNTPs, 1 mU/μL Pyrophosphatase, inorganic (0.1 U/μL) (Thermo Fisher Scientific), 1 U/μL RNasin Plus Ribonuclease Inhibitor (40 U/μL) (Promega), 0.6 µg MarathonRT and incubated for 1 h at 42 °C. The reactions were transferred to ice and ddH_2_O was added to yield 80 μL final volume. To detect the pyrophosphate (PPi) released during the reverse transcription, 20 µL of the diluted sample was further diluted with 60 µL of ddH_2_O (in triplicate) on a microplate, and 20 μL of DS solution was added. The samples were incubated 30 min at RT and the absorbance was measured at 620 nm using a Multiskan FC Microplate Photometer (Thermo Fisher Scientific).

### MarathonRT reaction optimization and storability

To estimate the optimal MRT/MRT-CBD units per 100 ng of bulk tRNA, 0.125 U, 0.25 U, 0.5 U, 0.75 U, 1 U, 1.25 U, 1.5 U and 1.75 units of MRT/MRT-CBD were used in RT reaction as described below. The cDNA was separated on denaturing electrophoresis on 6 % PAA/7 M urea/0.5× TBE gel. The best unit-to-template ratio was estimated from gel by densitometry analysis in ImageLab version 6.0 using the volumetric box function and relative cDNA amounts calculated in Excel.

The storability of MRT/MRT-CBD was assessed by preforming the activity assay (as described above) over the course of 52 weeks. For each measurement, the enzyme activity was evaluated either using a fresh aliquot that had not been thawed, using the same stock tube repeatedly to assess the effect of freeze-thaw cycles, or using a diluted stock.

### Total RNA extraction from yeast

*S. cerevisiae* BY4741 was grown at 30 °C, 200 rpm in 4× 250 mL YPD. The cells were harvested at OD_600_ ∼ 0.8 by centrifugation at 4,000 g for 10 min at 4 °C. The cell pellets were washed with 10 mL ice-cold 1× PBS (137 mM NaCl, 2.7 mM KCl, 10 mM Na_2_HPO_4_, 1.8 mM KH_2_PO_4_), re-suspended in 10 mL acidic phenol (pH 4.3, Carl Roth or Sigma) and stored at −20 °C. The next day, the pellets were melted at RT in a fume hood. Once melted, 10 mL of 0.9 % NaCl and 2 mL of 1-bromo-3-chloropropane (BCP) were added to the pellets. The cells were lysed by vortexing with glass beads and phase separation was performed by centrifugation at 10,000 g for 15 min at RT. The aqueous phase was transferred to a new tube containing 5 mL of acidic phenol and 1 mL of BCP, vortexed and centrifuged at 10,000 g for 10 min at RT. The aqueous phase was transferred to a new tube and combined with 2.5 vol. of 99.6 % EtOH. The total RNA was precipitated O/N at −20 °C and pelleted by centrifugation at 10,000 g for 30 min at 4 °C. Each pellet was washed 2× with 80 % EtOH with centrifugation at 10,000 g for 20 min at 4 °C after every wash. The pellets were air-dried and dissolved in 200 µL ddH_2_O. The quality and purity of total RNA were assessed by denaturing electrophoresis on 10 % PAA/7 M urea/0.5× TBE gel.

### Sequencing library preparation

The sequencing library was prepared following the mim-tRNAseq protocol^31^ with the following modifications.

#### tRNA deacylation and dephosphorylation

In a single reaction, 10 µg of the total RNA from *S. cerevisiae* was incubated with 1.5 µL 1 M Tris-HCl, pH 9 in a final volume of 21.5 µL at 37 °C for 45 min. No RNase inhibitor was added at this step, as the available RNase inhibitors are inactive at pH 9. The deacylation reaction was combined with 0.1U/µL T4 PNK (Thermo Fisher Scientific), 1× T4 PNK buffer A, 1 U/μL RNasin Plus Ribonuclease Inhibitor in the final volume of 100 µL and incubated at 37 °C for 45 min to dephosphorylate the tRNA fragments followed by purification on a column or gel.

#### A. tRNA spin column-based extraction

The deacylated and dephosphorylated total RNA was purified on NEB Monarch RNA Cleanup Columns (10µg) (T2037) or Macherey-Nagel NucleoSpin RNA XS Columns (740902.50S) following the published protocol^24^, and 16,000 g was used for all centrifugation steps as recommended by the column’s manufacturer. The samples were eluted in 10 µL 50 °C ddH_2_O. For the final library workflow 12 extractions were pooled, RNA concentration measured by NanoDrop, and purity assessed by denaturing electrophoresis on 8 % PAA/7 M urea/0.5× TBE gel.

#### B. tRNA gel-based extraction

The deacylated and dephosphorylated total RNA was resolved on 8 % PAA/7M urea/0.5× TBE gel alongside Low Range ssRNA marker (NEB) and visualized with SYBR Gold (Thermo Fischer Scientific). Species migrating at the size range of mature tRNAs (60 – 100 nt) were excised and gel slices crushed with disposable pestles in a microfuge tube. Following the addition of RNA gel elution buffer (300 mM sodium acetate pH 5.5, 1 mM EDTA) in a quantity enough to cover the gel pieces, the gel slurry was incubated on an end-over-end rotator at 4 °C O/N. Gel pieces were removed with Costar Spin-X centrifuge tube filters and RNA was recovered from the flow-through by adding an equal volume of isopropanol and precipitated at −70 °C for at least 1 h. The precipitated RNA was pelleted at 17,000 g for 1 h at 4 °C, washed twice with 1 mL ice-cold 80 % EtOH with centrifugation in between washes at 17,000 g for 20 min at 4 °C. The pellet was dried at 50 °C for 5 min and dissolved in 10 µL ddH_2_O. For the final library, 12 preparations were pooled, the concentration was measured by NanoDrop and purity was assessed by denaturing electrophoresis on 8 % PAA/7 M urea/0.5× TBE gel.

#### 3’ adapter pre-adenylation

The adapter pre-adenylation was done following the McGlincy and Ingolia protocol^45^.

#### 3’ adapter ligation

The column-extracted and gel-extracted tRNA was ligated to in-house pre-adenylated barcoded 3’adapters I1-I6 following the protocol from Behrens and Nedialkova^31^, except 1 U/μL RNasin Plus Ribonuclease Inhibitor (40 U/μL) was used in the 20 μL ligation reaction. The ligation products were separated on 8 % PAA/7 M urea/0.5× TBE gel and visualized with SYBR Gold. Species migrating at the size range of adapter-ligated tRNAs were excised and gel slices crushed with disposable pestles in a microfuge tube. Following addition of RNA gel elution buffer (300 mM sodium acetate pH 5.5, 1 mM EDTA) enough to cover the gel pieces, the gel slurry was incubated on an end-over-end rotator at 4 °C overnight. Gel pieces were removed with Costar Spin-X centrifuge tube filters and RNA was recovered from the flow-through by adding an equal volume of isopropanol and 2 µL of GlycoBlue (Thermo Fisher Scientific), vortexed thoroughly and precipitated at −70 °C for at least 1 h. The precipitated RNA was pelleted at 17,000 g for 1 h at 4 °C, washed twice with 1 mL ice-cold 80 % EtOH with centrifugation in between washes at 17,000 g for 20 min at 4 °C. The pellet was dried at 50 °C for 5 min and dissolved in 14 µL ddH_2_O per barcode and isolation type. The 3’adapter-ligated RNA was pooled by barcode and isolation type. Typical concentrations for the adapter-ligated tRNA were 80 – 106 ng/µL.

#### cDNA synthesis

Each reverse transcriptase had slightly different reaction mixes and conditions:

##### MRT/MRT-CBD

The reverse transcription reaction was performed in 20 µL final volume, where 100 ng of 3’adapter-ligated tRNA and 0.125 µM reverse transcription primer (5’-pRNAGATCGGAAGAGCGTCGTGTAGGGAAAGAG-iSp18-GTGACTGGAGTTCAGACGTGTGCTC-3’^31^ (Metabion) were mixed, denatured at 95 °C for 30 s, annealed for 5 min at RT and combined on ice with 0.5 mM dNTPs, 1 U/μL RNasin Plus Ribonuclease Inhibitor (40 U/μL), 0.5 U MarathonRT in 1× MRT reaction buffer (50 mM Tris pH 8.3, 200 mM KCl, 2.5 mM MgCl_2_, 5 mM DTT, 20 % glycerol) and incubated for 16 h at 42 °C.

##### uMRT

The reverse transcription reaction was performed in 20 µL final volume, where 100 ng of 3’adapter-ligated tRNA, 0.1 µM reverse transcription primer (as noted above) and 0.5 mM dNTPs were mixed, denatured at 95 °C for 30 s, snap-cooled on ice for 2 min and combined on ice with 1 U/μL RNasin Plus Ribonuclease Inhibitor (40 U/μL), 1 U/μL uMRT (20 U/μL, RNAConnect) in 1× uMRT reaction buffer and incubated for 16 h at 42 °C.

##### Induro

The reverse transcription reaction was performed in 20 µL of final volume, where 100 ng of 3’adapter-ligated tRNA and 0.5 µM reverse transcription primer (as noted above) were mixed, denatured at 82 °C for 2 min, annealed on RT for 5 min and combined on ice with 10 mM DTT, 1 U/μL RNasin Plus Ribonuclease Inhibitor (40 U/μL), 10 U/μL Induro (200 U/μL, NEB) in 1× Induro RT reaction buffer and incubated at 42 °C for 10 min. After the incubation, 1.5 μL of 10 mM dNTPs were added to the reaction and incubated for 16 h at 42 °C (Nakano et al., 2025).

After cDNA synthesis, the RNA was hydrolysed by adding 1 μL of 5 M NaOH, heated at 95 °C for 3 min and transferred to ice. The cDNA was separated immediately on 6 % PAA/7M urea/0.5× TBE gel alongside Low Range ssRNA marker and visualized with SYBR Gold. Species migrating at the size range of cDNA were excised and gel slices crushed with disposable pestles in a microfuge tube. Following the addition of DNA gel elution buffer (10 mM Tris pH 8, 300 mM NaCl, 1 mM EDTA), the gel slurry was incubated on an end-over-end rotator at 4 °C O/N. Gel pieces were removed with Costar Spin-X centrifuge tube filters and cDNA was recovered from the flow-through by adding an equal volume of isopropanol, 2 µL of GlycoBlue and precipitating at −70 °C for at least 1 h. The precipitated cDNA was pelleted at 17,000 g for 1 h at 4 °C, washed twice with 1 mL ice-cold 80 % EtOH with centrifugation in between washes at 17,000 g for 20 min at 4 °C. The pellet was dried at 50 °C for 5 min and dissolved in 5.5 µL ddH_2_O per pooled RT reaction. cDNA quality was assessed on TapeStation (Agilent) using HS RNA ScreenTape.

### cDNA circularization and library construction PCR

The cDNA was circularized using CircLigase (Lucigen) and 5.5 µL of the pooled cDNA containing all the reverse transcription replicates per enzyme and isolation type. The circularized cDNA was directly used for library construction PCR using KapaHiFi (Roche) and 4 amplification cycles. The PCR products were purified using Select-a-Size DNA Clean & Concentrator MagBead Kit (Zymo) following the protocol for remaining fragment size >100 bp by the manufacturer using 1.5 mL microfuge tubes and eluting in 12 µL with room temperature ddH_2_O. The purified DNA concentration was measured using Qubit dsDNA HS kit (Thermo Fischer Scientific) on a fluorometer. The libraries were pooled equimolarly and sequenced for 300 cycles on NovaSeqX 10 B XP flowcell (PE, 150 bp, Illumina).

### Sequencing data analysis

Only R1 reads were used for downstream analysis. R1 reads were processed and analysed using mim-tRNAseq toolkit (v1.3.9) following the general workflow described by Behrens and Nedialkova^31^ and the mim-tRNAseq documentation. mim-tRNAseq toolkit was executed with the following non-default parameters: --no-cca, --cluster-id 0.90, --min-cov 0.0005, --max-mismatches 0.1, --max-multi 4, and --remap. Remapping was enabled with a mismatch tolerance of 0.075. Reads were mapped to *S. cerevisiae* nuclear and mitochondrial tRNA reference sequences provided with the mim-tRNAseq package. A reference set excluding *E. coli* spike-in tRNAs was used, as no spike-in was included during library preparation. Reference files were supplied explicitly to mim-tRNAseq using the -t, -o, and -m parameters. Alignment and clustering were performed using GMAP (v2019.02.26). Coverage plots, read counts and mismatch profiles were generated using mim-tRNAseq’s built-in plotting functions, except mapping statistics, where the raw data was used to replot in Excel. When replicate data were summarized, values are reported as mean ± standard deviation. Correlation analyses between replicates were quantified using Pearson correlation coefficients.

### Liquid chromatography mass spectrometry analysis

Sample preparation and analysis were performed as described previously^46^. The data analysis was performed using Mzmine version 4.5.37. The results were normalized using internal standard 1,3-dimethylpseudouridine. The data was visualized in R version 2025.05.1.

## Acknowledgments

The authors thank Salla Kalaniemi and Sari Korhonen for their valuable technical assistance. The authors wish to acknowledge Dr. Nina Sipari, Helsinki Metabolomics Centre (formerly Viikki Metabolomics Unit) for mass spectrometry analysis, the Next Generation Sequencing Facility at Vienna BioCenter Core Facilities (VBCF), member of the Vienna BioCenter (VBC), Austria, for tRNA sequencing, the HiLIFE Biocomplex Unit, University of Helsinki—a member of Instruct-ERIC Centre Finland, FINStruct, and Biocenter Finland—for high-speed and ultracentrifugation services. All schematic representations created with https://BioRender.com and modified in Inkscape version 1.4.3. Finally, the authors thank all members of the RNAcious laboratory for their insightful intellectual discussions regarding this work.

## Author contributions

**Jenni Katri Pedor:** Conceptualization, Methodology, Validation, Formal analysis, Investigation, Data Curation, Writing - Original Draft, Writing - Review & Editing, Visualization **Pavlina Gregorova:** Methodology, Formal analysis, Data Curation, Writing - Review & Editing **Matea Radesic:** Formal analysis, Investigation, Data Curation, Writing - Review & Editing, Visualization **L. Peter Sarin:** Conceptualization, Resources, Supervision, Project administration, Funding acquisition, Writing - Review & Editing.

## Data availability statement

The data that support this study are available from the corresponding author upon reasonable request. All MRT-tRNA-seq raw files generated during this study are available for download on the NCBI Sequence Read Archive (SRA) database under BioProject accession number PRJNA1427126.

## Supporting information

This article contains supporting information^16,17,31^.

## Funding and additional information

This work was supported by the Novo Nordisk Foundation [grant no. NNF19OC0054454 to L.P.S.], the Research Council of Finland [grant no. 354906 to L.P.S.], and the Sigrid Jusélius Foundation [grant no. 240192 to L.P.S.]. J.K.P., P.G., and M.R. are fellows of the Doctoral Programme in Integrative Life Science, University of Helsinki. Any opinions, findings, and conclusions or recommendations expressed in this material are those of the authors and do not necessarily reflect the views of the funding bodies.

