## Supplementary Information for "Production of MarathonRT and its comparison with commercial reverse transcriptases for tRNA sequencing library preparation"

**Contents:**

Supplementary Results S1-S2

Supplementary Figures S1-S9

Supplementary File S1 (XLSX) – Densitometry analysis from WB.

Supplementary File S2 (XLSX) – Raw data and analysis from Bradford Assay.

Supplementary File S3 (XLSX) – Raw data and analysis of the enzymes' activity for 52 weeks.

Supplementary File S4 (XLSX) – Enzymes' cost-per-reaction calculations.

Supplementary File S5 (XLSX) – Raw data and analysis of the enzymes' freeze-thaw endurance.

Supplementary File S6 (XLSX) – Raw data and analysis of the diluted enzymes' storability.

Supplementary File S7 (XLSX) – Densitometry analysis of optimal enzyme-unit-to-template ratio.

Supplementary File S8 (XLSX) – Sequencing data general and mapping statistics.

Supplementary File S9 (XLSX) – Identified tRNA isoacceptors.

Supplementary File S10 (XLSX) – Ions identified by LC-MS analysis.

**Supplementary Results**

Supplementary Result 1 – extraction-dependant presence of 5S rRNA in tRNA samples

We noticed a 5S rRNA presence in the final tRNA samples (Figure 2D). This was surprising given that prior results with the column-based method did not contain trace amounts of 5S rRNA (Figure 2C), whereas gel extraction was performed by carefully excising the tRNA containing region. It is conceivable that a slight overload of the gel could have an adverse impact on separation, thus accounting for the trace amounts of rRNA detected in the gel extracted sample. Nonetheless, the column-based enrichment method was performed using the same input amounts as above (2.5-10 µg of total RNA)—yet 5S rRNA was present in all samples (Supp. Fig. S3C). It should be noted that the only variable that changed between these samples was the acidic phenol used for extraction. Indeed, we observed that acidic phenol sourced from Sigma enriches short RNAs whereas the equivalent product from Carl Roth is less biased towards short RNAs (Supp. Fig. S3D, E). This highlights the fact that seemingly identical products may have different properties depending on the manufacturer.

Supplementary Result 2 – sequencing library preparation following mim-tRNAseq protocol[30]

The pre-adenylated indexed 3'adapter was ligated to the tRNA extracted either by spin column or gel extraction (Supp. Fig. S4). The gel-purified, adapter-ligated tRNA was used as a template for the reverse transcription reactions following the manufacturer's conditions for each enzyme, except for prolonging the reaction time to 16 h for all reactions, as it has been reported to enhance the production of full-length cDNA[16,17] (Supp. Fig. S5). Each RT reaction was performed in 6 technical duplicates. The cDNA was circularized by CircLigase followed by sequencing library amplification using KapaHiFi and Illumina RPI sequencing primers. The quality of each library after purification was checked on TapeStation (Agilent) HS DNA tapes (Supp. Fig. S6, A-H) followed by equimolar sequencing library preparation (Supp. Fig. S6, I).

**Supplementary Figures**

A.

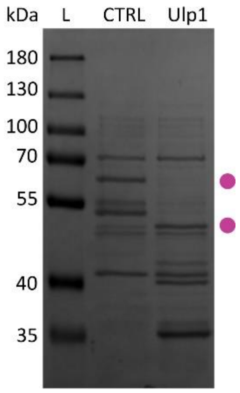


**Supplementary Figure S1. C-terminally truncated MRT present when produced in *E. coli* but not in insect cells.** Bands representing full-length MRT are indicated by a dot. Marker: PageRuler™ Prestained Protein Ladder, 10–180 kDa (Thermo Scientific). (A) SUMO-tag cleavage by Ulp1 analysed on 10 % SDS-PAGE. *CTRL* – affinity purified MRT; *Ulp1* – SUMO-tag cleavage reaction.

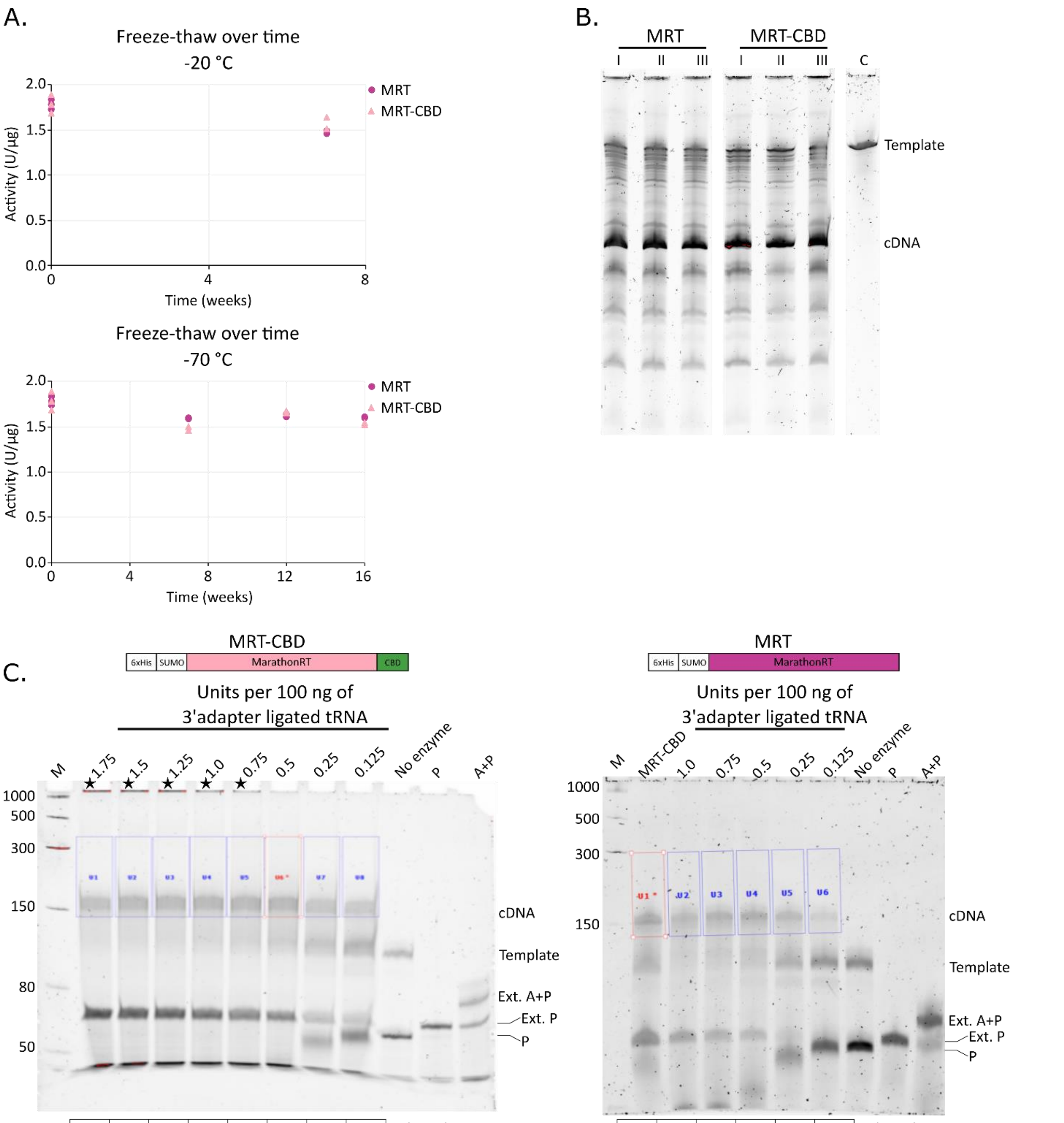


| V1 | V2 | V3 | V4 | V5 | V6 | V7 | V8 | Volume box |
|---|---|---|---|---|---|---|---|---|
| 0.778 | 0.902 | 0.899 | 0.975 | 0.993 | 1 | 0.682 | 0.682 | Relative quantity |
| 1.75 | 1.5 | 1.25 | 1 | 0.75 | 0.5 | 0.25 | 0.125 | Units |

| V1 | V2 | V3 | V4 | V5 | V6 | Volume box |
|---|---|---|---|---|---|---|
| 1 | 0.773 | 0.937 | 1.022 | 0.676 | 0.255 | Relative quantity |
| 0.5* | 1 | 0.75 | 0.5 | 0.25 | 0.125 | Units |

**Supplementary Figure S2. MRT and MRT-CBD activity and enzyme-to-tRNA ratio.** (A) Activity of diluted protein samples (0.6 mg/mL) after each freeze-thaw cycle over 8 and 16 weeks after storage at −20 °C and −70 °C, respectively. Measurements were performed in technical duplicates, except the first time point (n = 3). (B) Analysis of the RT reaction used for activity assay on 5 % denaturing urea-PAA gels, loaded 1/12th of reaction. *I–III* – replicate reactions; *C* – no-enzyme control; *Template – in vitro* produced RNA (642 nt); *cDNA* – 277 nt. (C) RT reaction analysis on 6 % denaturing urea-PAA gels for densitometry quantification of free cDNA using ImageLab v 6.0 volume box feature. Controls: *No enzyme* – no enzyme added, *P* – no template added, RT primer only, *A + P* – no template added, 3′-adapter + primer. *cDNA* – 150–164 nt, *Template* – 3′-adapter-ligated tRNA, 111–125 nt, *Ext. A+P* – extended 3′-adapter:RT-primer duplex, *Ext. P* – extended primer. Left panel: MRT-CBD (star symbol indicates RNA/cDNA bound to enzyme, visible in well). Right panel: MRT.

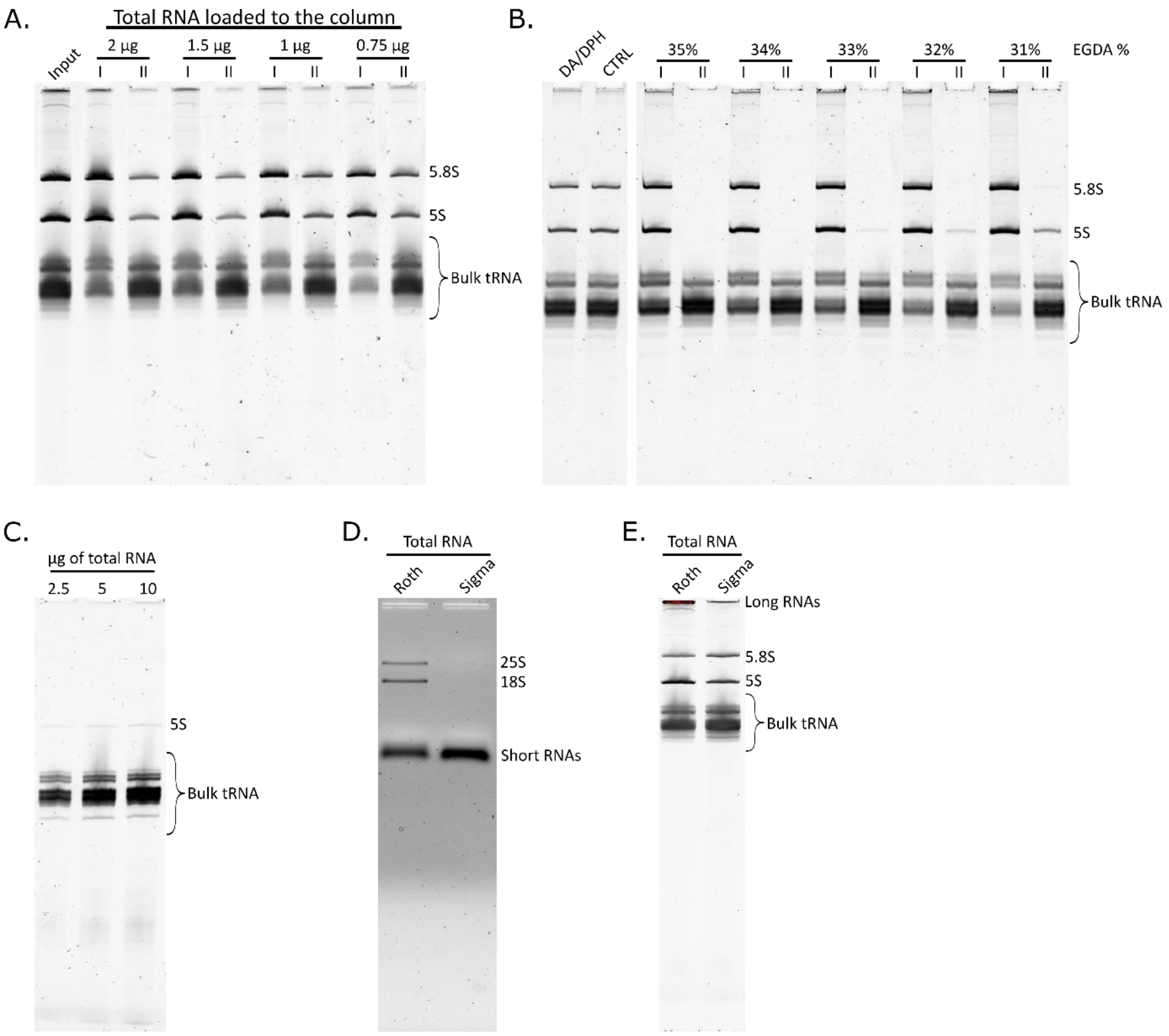


**Supplementary Figure S3. Silica spin column-based tRNA extraction optimization.** Bands corresponding to 5.8S, 5S, 18S, and 25S rRNA indicated. (A) RNA eluted from NucleoSpin RNA XS Columns (Macherey-Nagel) on 10 % denaturing urea-PAA gel. *Input* – total RNA post-phenol/BCP purification; *I/II* – eluates from column I and II. (B) Column cut-off dependence on EGDA concentration. *DA/DPH* – deacylated and dephosphorylated total RNA; *CTRL* – only dephosphorylated total RNA; *I/II* – eluates from column I and II. (C) Total RNA (2.5, 5, 10 µg) extracted by Roth phenol loaded to Monarch RNA Cleanup Columns (10 µg, NEB); eluates from column II analysed on 8 % denaturing urea-PAA. (D, E) Total RNA extracted with Roth or Sigma phenol analysed on 1.5 % agarose gel (D) and 10 % denaturing urea-PAA (E).

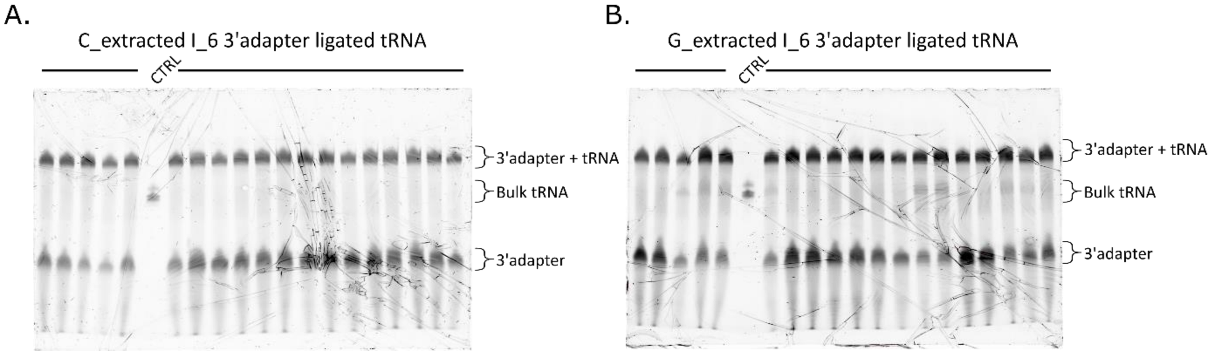


**Supplementary Figure S4. Representative reactions of I6-indexed-3'adapter ligation to column and gel extracted tRNA analysed on 8 % denaturing urea-PAA.** *CTRL* – bulk tRNA. Band identities indicated next to gels. (A) On-column extracted tRNA ligation reactions. (B) Gel extracted tRNA ligation reactions.

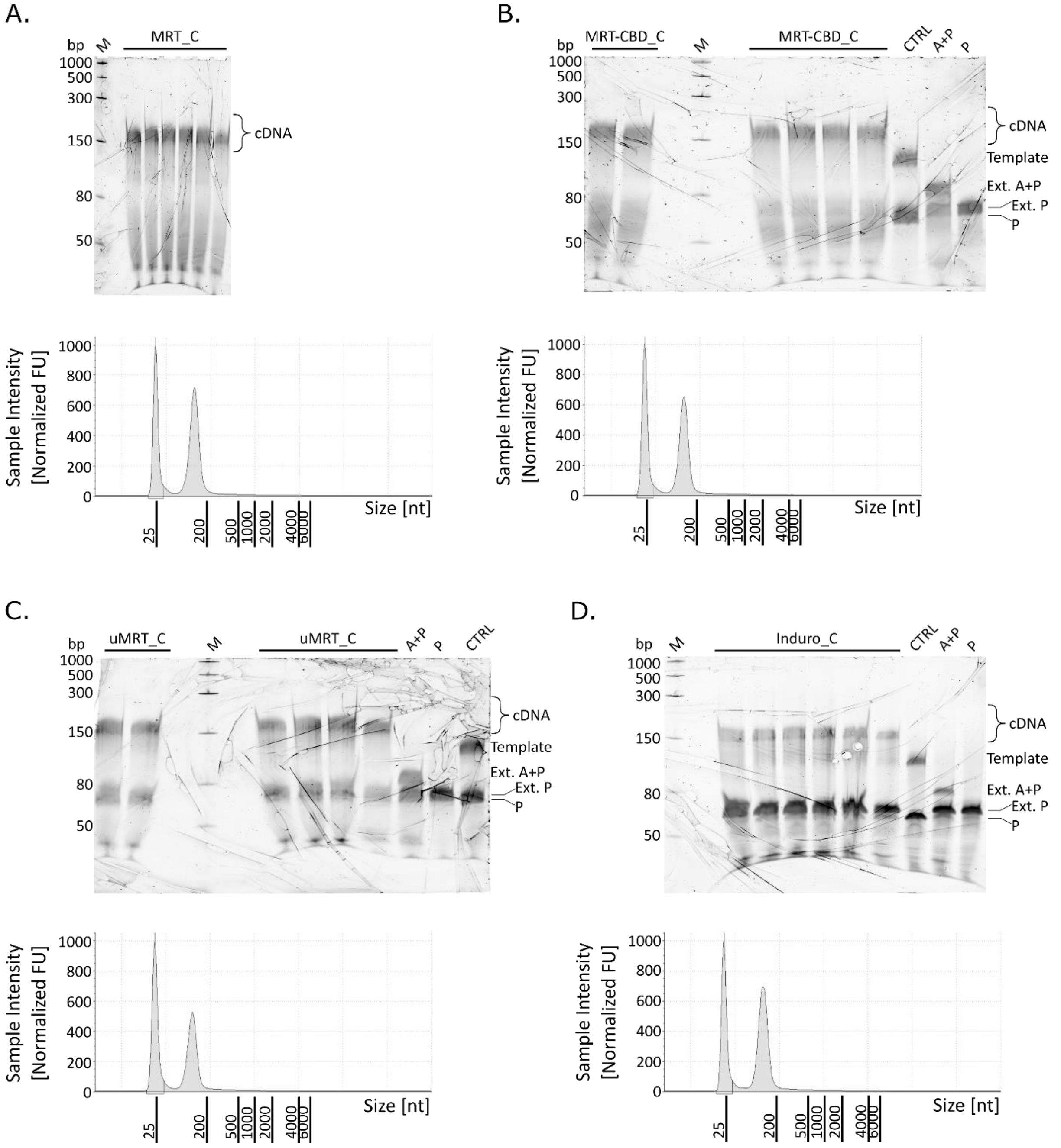


**Supplementary Figure S5. RT reactions on 6 % denaturing urea-PAA and gel extracted cDNA analysed via TapeStation.** Representative results using column (_C) extracted tRNA. Top panels: urea-PAA gels. Marker: Low Range ssRNA (NEB). Controls: *CTRL* – reaction with no enzyme added; *P* – no template added, RT primer only; *A + P* – no template added, 3′-adapter + primer. *cDNA* – 150–164 nt; *Template* – 3′-adapter-ligated tRNA, 111–125 nt; *Ext. A+P* – extended 3′-adapter:RT-primer duplex, *Ext. P* – extended RT primer. Lower panels: TapeStation profiles. The 25 nt peak that is present in all samples corresponds to the loading size marker. (A) MRT. (B) MRT-CBD. (C) uMRT (RNAConnect). (D) Induro (NEB).

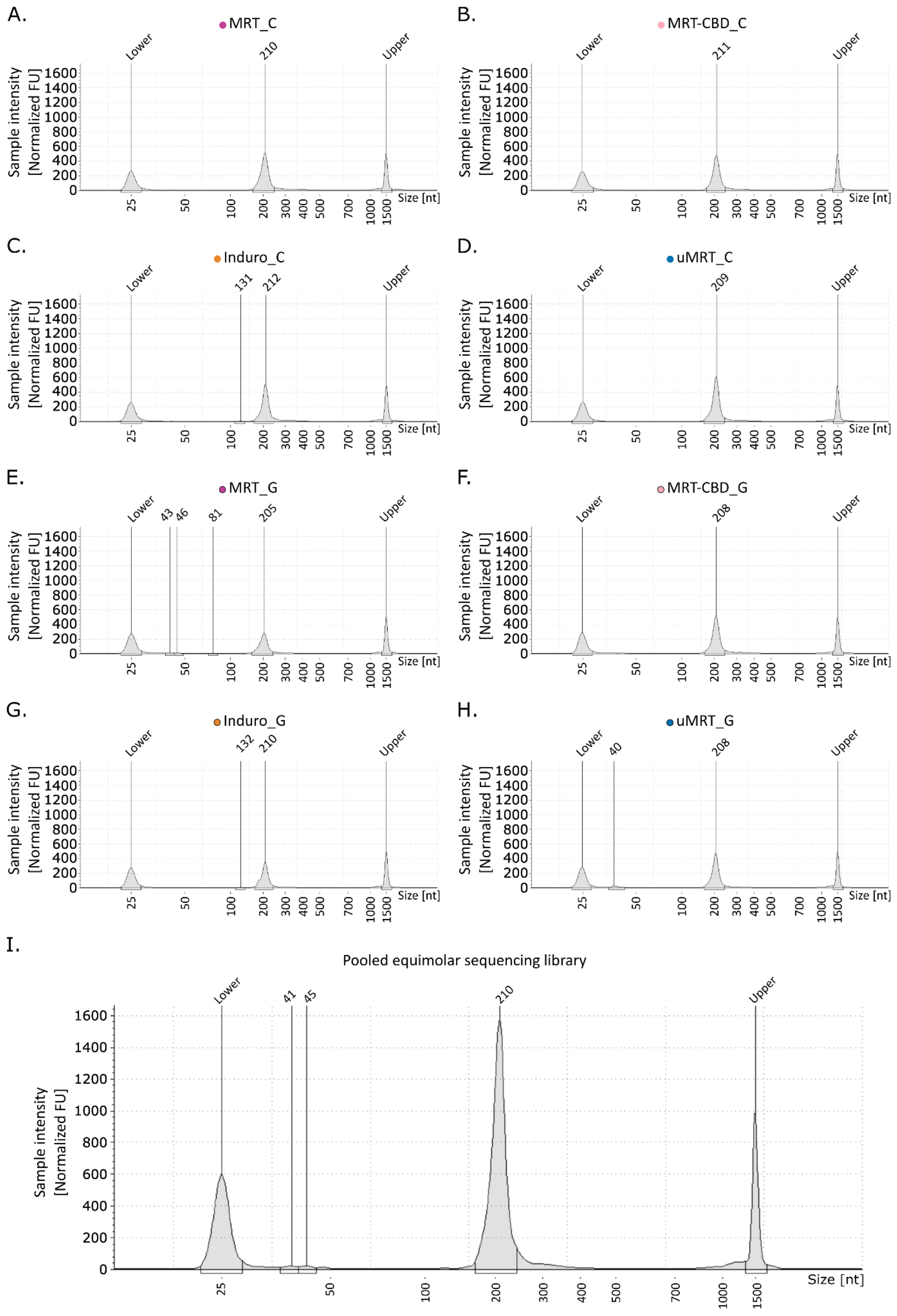

A.
MRT_C
B.
MRT-CBD_C
C.
Induro_C
D.
uMRT_C
E.
MRT_G
F.
MRT-CBD_G
G.
Induro_G
H.
uMRT_G
I.
Pooled equimolar sequencing library
Lower
Upper
210
211
131
212
209
43
46
81
205
208
132
40
41
45
Sample intensity [Normalized FU]
0
200
400
600
800
1000
1200
1400
1600
Size [nt]
25
50
100
200
300
400
500
700
1000
1500

**Supplementary Figure S6. Sequencing libraries analysed via TapeStation.** The 25 nt and 1500 nt peak that are present in all samples correspond to the loading size marker. (A-H) Sequencing libraries per enzyme and extraction type, pooled 6 technical replicates. (I) Pooled equimolar library sent to sequencing.

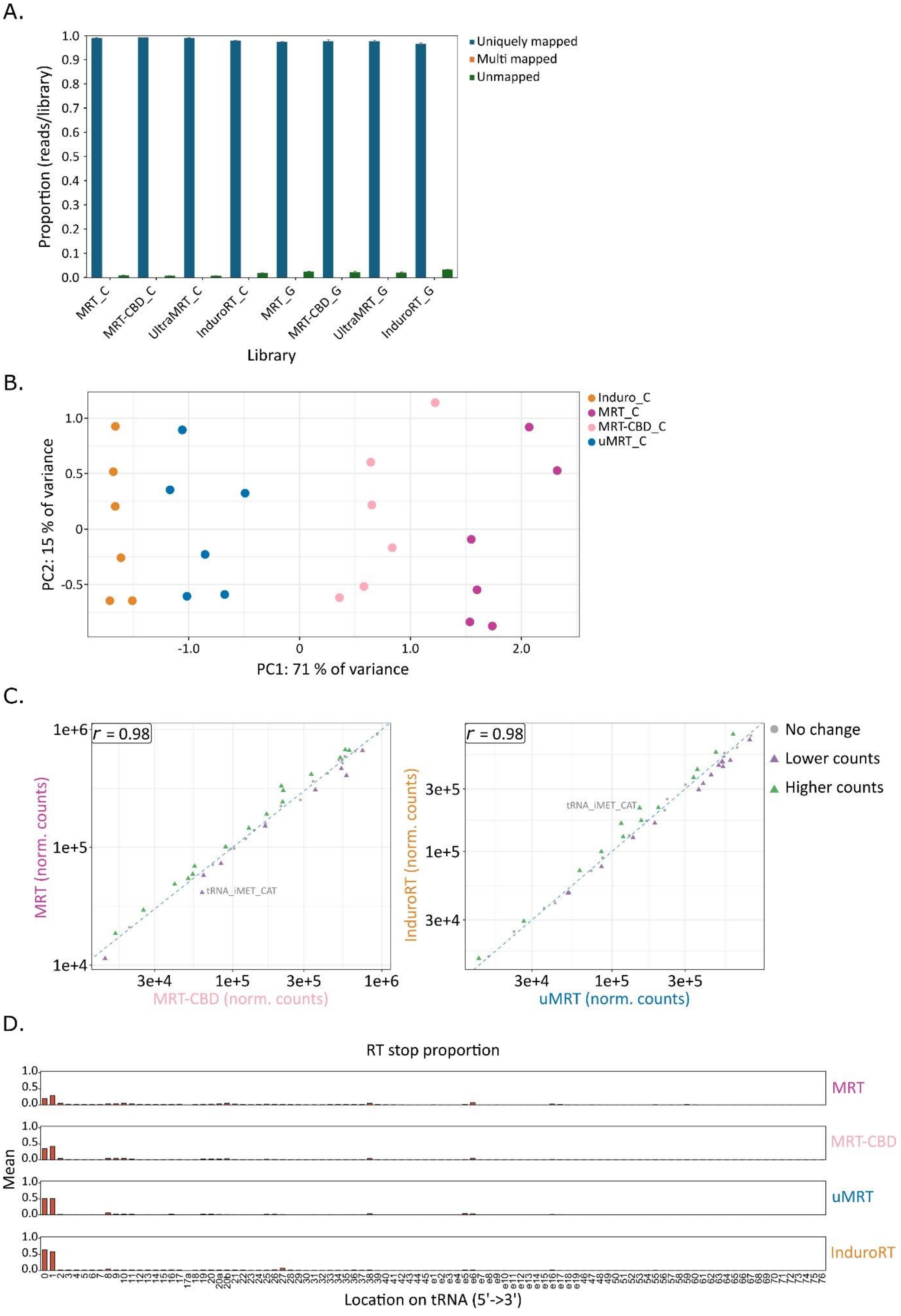

A.
Uniquely mapped
Multi mapped
Unmapped
Proportion (reads/library)
MRT_C
MRT-CBD_C
UltraMRT_C
InduroRT_C
MRT_G
MRT-CBD_G
UltraMRT_G
InduroRT_G
Library
B.
Induro_C
MRT_C
MRT-CBD_C
uMRT_C
PC2: 15 % of variance
PC1: 71 % of variance
C.
r = 0.98
MRT (norm. counts)
MRT-CBD (norm. counts)
tRNA_iMET_CAT
r = 0.98
InduroRT (norm. counts)
uMRT (norm. counts)
tRNA_iMET_CAT
No change
Lower counts
Higher counts
D.
RT stop proportion
Mean
MRT
MRT-CBD
uMRT
InduroRT
Location on tRNA (5'->3')

**Supplementary Figure S7. Mapping statistics, sample clustering, tRNA count comparison and accumulative RT stops.** (A) Sequencing mapping statistics showing fraction of reads uniquely mapped, multi-mapped, and unmapped per library (mean ± SD, n = 6). (B) PCA of column extracted tRNA samples (n = 24) coloured by RT enzyme. Each RT reaction performed in six technical replicates. (C) Scatter plots comparing average normalized tRNA counts across MRT/MRT-CBD and InduroRT/uMRT (technical replicates n = 6); Axes represent average normalized read counts from DESeq2. Pearson's *r* indicated. (D) Mean position-specific RT stop frequencies aggregated over technical replicates (n = 6) per enzyme (coverage threshold >0.0005).

A.

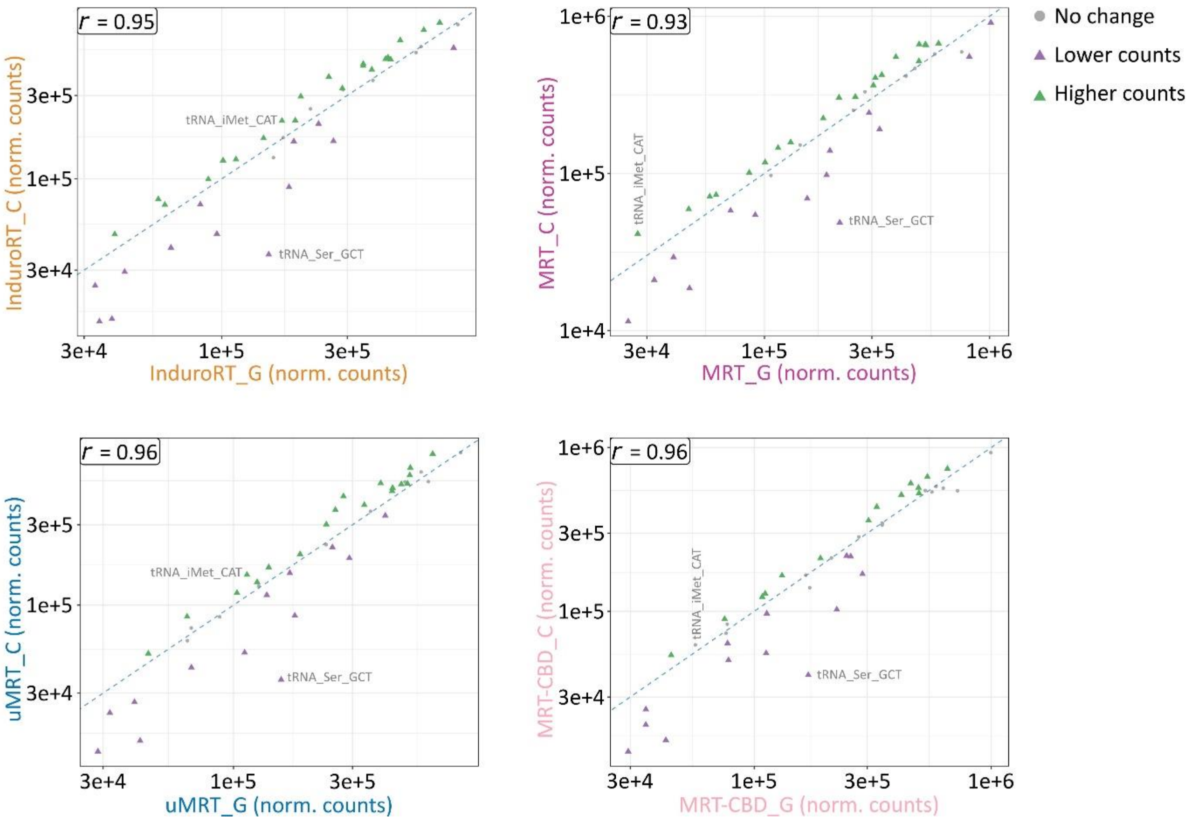


**Supplementary Figure S8. Correlation of tRNA counts between column and gel extracted samples.** (A) Read counts per tRNA extraction type (n = 6) are compared across enzymes. Axes represent average normalized read counts from DESeq2. Pearson (*r*) indicated.

A.

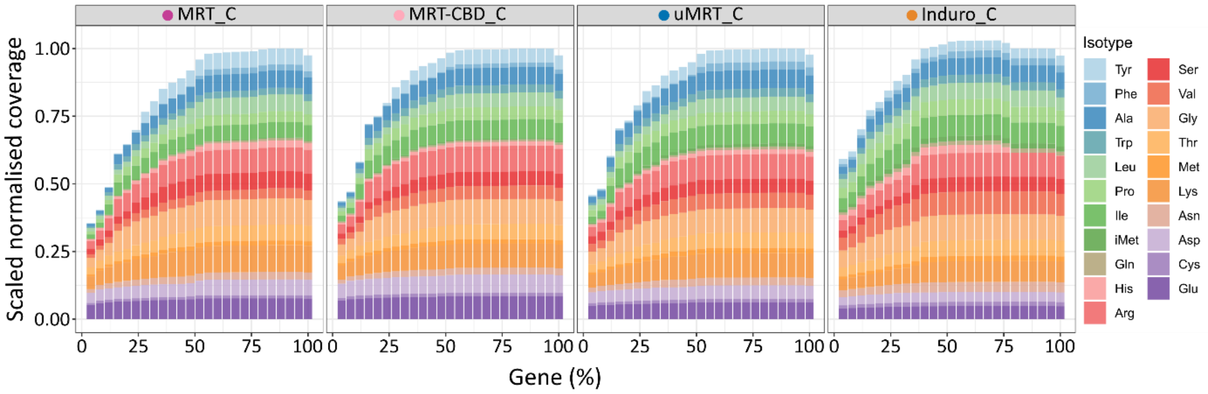


**Supplementary Figure S9. Representative tRNA coverage profiles per RT enzyme.** Metagene plots showing normalized read coverage along nuclear-encoded mature tRNA isotypes (5′→3′) after removal of terminal CCA. Coverage averaged across all detected tRNA species for each enzyme per representative sample.
